# Biochemical, structural, and functional characterization of the *Nocardia asteroides* dihydrofolate reductase: a primary target of anti-nocardiosis treatment

**DOI:** 10.64898/2026.05.27.728178

**Authors:** Julie Couston, Sébastien Lainé, Jérôme Feuillard, Mickaël Blaise

## Abstract

Nocardiosis is a human infectious disease caused by several species of *Nocardia* and primarily affecting the skin, lungs and central nervous system. The first line treatment is based on cotrimoxazole, combining trimethoprim and sulfamethoxazole. These two drugs target respectively the dihydrofolate synthase (DHFR) and the dihydropteroate synthase (DHPS) involved in the essential folate synthesis pathway. The occurrence of drug resistance to these two drugs is however frequent. While the molecular mechanisms of trimethoprim resistance are well documented in other bacteria, they remain poorly explored and documented in *Nocardia*. This is partly because few biochemical structural or genetic studies have been conducted on DHFR from this genus. In this study, we report the biochemical and structural characterization of DHFR from *Nocardia asteroides* (DHFR_Nad_). We show that overexpression of DHFR_Nad_ in *N. asteroides* confers strong resistance to trimethoprim. We recombinantly expressed and purified active DHFR_Nad_ and determined its inhibition constant for trimethoprim. We solved the crystal structure of DHFR_Nad_ bound to trimethoprim at high resolution. Further, biochemical studies of mutant DHFR variants pinpointed the role of important residues for trimethoprim binding and drug-resistance.

**Highlights:** First biochemical and structural characterization of *Nocardia asteroides* DHFR.

Overexpression of DHFR_Nad_ induces high-level trimethoprim resistance in *N. asteroides*.

Crystal structure of DHFR_Nad_ reveals key residues for trimethoprim binding.

Mutagenesis confirms residues critical for trimethoprim susceptibility.

IC_50_ data confirm strong DHFR_Nad_ inhibition by trimethoprim and methotrexate

## Introduction

*Nocardia asteroides* is a gram-positive bacterium belonging to the actinobacterial phylum and corynebacteriales/mycobacteriales order. *Nocardia* species (*Nocardia* spp.) are environmental bacteria that can cause mild to severe human infections named nocardiosis. *N. asteroides* is one of these pathogenic *Nocardia* spp. able to infect the skin, the lungs and the brain [1]. Albeit cutaneous infections can usually be treated, pulmonary and central nervous system (CNS) infections are particularly severe, difficult to treat and life-threatening [2,3].

The current recommended regimen to treat nocardiosis is a combination of sulfamethoxazole (SMX) and trimethoprim (TMP) [4] targeting two enzymes of the folate pathway, namely the dihydropteroate synthase and the dihydrofolate reductase (DHFR) respectively [5]. DHFR is a ubiquitous and essential enzyme for living organisms that catalyzes the reduction of dihydrofolate to tetrahydrofolate and using NADPH as a cofactor [6]. The anti-nocardiosis treatment based on cotrimoxazole is efficient but since it is given for a long period of time, weeks to months, resistance to one or both molecules is often reported [1,7–9]. Mehta and collaborators have demonstrated experimentally that exposure of *Nocardia* strains to TMP-SMX triggers resistance and demonstrated notably that TMP resistance was linked to mutations affecting the gene encoding for the DHFR. Further structural modelling approaches suggested that these mutations may affect residues that are part of or near the putative active site of the DHFR [10].

DHFR has been well characterized by genetic, biochemical and structural approaches in numerous bacteria. In *E. coli* the impact of DHFR mutations in relation to TMP resistance has been thoroughly investigated [11–13]. However, to date and to our knowledge only one biochemical study from the 1980’s reported the direct inhibition of a DHFR that was isolated from the *Nocardia restricta 1430* species [14]. Further, there is no structural data on DHFR from any *Nocardia* species. In the absence of biochemical and structural data it may therefore be difficult to apprehend and fully understand the impact of either acquired or innate resistant mutations found in clinical strains.

In the scope of better characterizing *Nocardia* DHFR, we engaged in biochemical, structural and functional studies of DHFR (DHFR_Nad_) from *N. asteroides* NBRC 15531. We show that overexpression of DHFR_Nad_ in *N. asteroides* affects the susceptibility to trimethoprim. We report the purification of a recombinantly expressed and active DHFR_Nad_ as well as its inhibition constants for trimethoprim. We determined the crystal structure of DHFR_Nad_ at high resolution bound to trimethoprim which highlights residues previously found in *Nocardia* species resistant to this antibiotic.

## Materials and Methods

### 1- Gene amplification, cloning and mutagenesis for recombinant protein expression

The *folA* gene was amplified with Phusion polymerase from *N. asteroides* NBRC 15531 genomic DNA by PCR using the primers forward: 5’-agcagacggatcggcctgatctgggcgcagg-3’and reverse: 5’-tcagcggcgggtgtagtggcgaacgcgatatctcagg-3’. The PCR fragment was purified on agarose and 3’ overhang A nucleotide was added with the Taq Polymerase to the PCR product and cloned into Champion*™* pET SUMO (ThermoFisher SCIENTIFIC) in frame with the multi-histidine and SUMO tags. Correct cloning was assessed by restriction analysis and the final construct was verified by whole plasmid sequencing (Eurofins MWG). The final sequence encodes residues 2-161 of the U5E929 Uniprot protein.

The DHFR variants, D30A, M31A and F34A were generated by site-directed mutagenesis starting from the pET SUMO::*dhfr*_Nad_ construct. To do so, we used the Q5^®^ Site-Directed Mutagenesis Kit (NEW ENGLAND Biolabs) and the following primers: DHFRmut-D30A-Fw: 5’-ggcgcgtccccgaggCcatggccaacttccgcgacacca-3’; DHFRmut-D30A/M31A-Rv:5’-acggaatcgtgttgtccaccccgatgacgccg-3’; DHFRmut-M31A-Fw:5’-ggcgcgtccccgaggacGCggccaacttccgcgacacc-3’; DHFRmut-F34A-Fw: 5’-cccgaggacatggccaacGCccgcgacaccaccatgggcc-3’; DHFRmut-F34A-Rv: 5’-gacgcgccacggaatcgtgttgtccacccc-3’, with the mutations introduced highlighted by the nucleotides in capital letters. The reactions were essentially performed following the manufacturer instructions, with the exception of the PCR cycles that were performed as follows: denaturation step at 98 ℃ for 2 min; 30 cycles of : 98 ℃ for 10 s, 70 ℃ for 30 s, 72 ℃ for 4 min; and the final extension at 72 ℃ for 2 min. The mutants were further verified by sequencing.

### 2- Gene amplification and cloning for overexpression studies in *Nocardia asteroides*

The *dhfr*_Nad_ or mutant genes were amplified from the pET SUMO::*dhfr*_Nad_ or mutant constructs with the following primers forward: 5’ gtgcc<u>GAATTC</u>atgagcagacggatcggcctgatctgggcgcaggccgcgaac-3’ and Reverse: 5’-gtgcc<u>GGATCC</u>ttattaCTTCTCGAACTGCGGGTGGCTCCAgcggcgggtgtagtggcgaacgcgatatctcag gcccgatttcg-3’and cloned between the EcoRI and BamHI restriction sites of a modified pNV118 *Nocardia* plasmid derived from the original pNV18 [15]. The reverse primer harbors a Strep-Tag II sequence (capital letters) situated upstream the two stop codons and enabling the detection of the protein expression by western blot. Of note, the original pNV118 plasmid was modified so that the *lac* promoter was replaced by the promoter of the ampicillin resistance gene, and the kanamycin selection marker was exchanged to zeocin.

### 3- Transformation of *N. asteroides* by electroporation

A 50 mL culture of *N. asteroides* was grown exponentially in Brain Heart Infusion (BHI) media supplemented with 1 % glycine then spun down at 4000 *g* for 15 min. The bacterial pellet was resuspended in sterile and cold water and spun down at 4000 *g* for 10 min. This step was repeated twice. After the third wash in water, the bacteria were resuspended in 10 % sterile glycerol and centrifuged at 4000 *g* for 10 min. Bacteria were then resuspended in 2 mL of glycerol 10 % and passed through a G26 needle plugged to a tuberculin syringe 20 times. Then 1 μg of plasmid was incubated with 50 μL of bacteria for 30 min on ice and transferred into an electroporation cuvette (Gene Pulser cuvette, 0.2 cm Bio-rad). The bacteria were electroporated at 2.5 kV in a MicroPulser electroporator (Bio-Rad), transferred into an microtube with 250 μL of BHI media and grown for 3 h at 37℃ before being plated onto BHI agar media supplemented with 50 μg.mL^-1^ of Zeocin.

### 4- Expression and purification

The pET SUMO::*dhfr*_Nad_ construct was transformed into an *E. coli* BL21 strain resistant to phage T1 (New England Biolabs) carrying the pRARE2 plasmid (Novagen). Colonies were selected on Luria-Bertani (LB) agar plates supplemented with kanamycin (50 μg.mL^-1^) and chloramphenicol (30 μg.mL^-1^). A single colony was used to start a preculture in LB liquid media with the two antibiotics overnight (ON) at 37°C under agitation. The day after, the preculture was used to inoculate 12 L of the same media divided into 4 large flasks. The culture was grown under agitation at 37°C until the OD_600nm_ reached 0.8 and was placed into melting ice for 30 min. The protein expression was induced by adding 1 mM isopropyl-thio-galactoside and the culture was grown ON at 18°C under agitation. The bacterial pellet was collected after a 15 min centrifugation at 6000 *g*, then resuspended in lysis buffer: 50 mM Tris-HCl pH 8, 200 mM NaCl, 20 mM imidazole and 5 mM β-mercaptoethanol. The cells were lysed by sonication and the soluble protein extract was obtained after a centrifugation step at 27000 *g* for 45 min. The supernatant was loaded twice onto 3 gravity columns containing 2 mL of Ni-sepharose-6-fast-flow beads (Cytiva) each previously equilibrated with lysis buffer, for an immobilized-metal affinity chrommatography (IMAC) step. The columns were then washed with 8 mL of lysis buffer, then 8 mL of wash buffer: 50 mM Tris-HCl pH 8, 1 M NaCl and 5 mM β-mercaptoethanol. The protein was eluted with 10 mL of elution buffer: 50 mM Tris-HCl pH 8, 200 mM NaCl, 250 mM imidazole and 5 mM β-mercaptoethanol. The SUMO tag was cleaved ON with the SUMO protease (1/200 w/w ratio) at 4°C during dialysis against dialysis buffer: 50 mM Tris-HCl pH 8, 200 mM NaCl, 10 % glycerol and 5 mM β-mercaptoethanol. The protein was then loaded 3 times onto a gravity column made of 4 mL Ni-sepharose-6-fast-flow beads and equilibrated with dialysis buffer. The cleaved protein was recovered into the flow-through, concentrated to 5 mg.mL^-1^ and subjected to a last polishing step by size-exclusion chromatography (SEC) onto a Superdex 75 Increase 10/300 GL column (Cytiva) in SEC buffer: 20 mM Tris-HCl pH 8, 200 mM NaCl, 10 % glycerol and 1 mM β-mercaptoethanol for crystallization purpose. The purification protocol for wild-type (WT) and variant (D30A, M31A and F34A) DHFR used for activity assays was similar, with the exception that the last SEC step was performed in 50 mM Bis-Tris pH 5.5, 200 mM NaCl, 5 mM β-mercaptoethanol and 10 % glycerol. The purity of the WT and variants was estimated to be superior to 95 % as judged by Coomassie-stained SDS-PAGE.

### 5- Crystallization

The protein solution at 15 mg.mL^-1^ was supplemented with 2 mM of NAPDH (stock solution at 30 mM diluted in water) and 2 mM of trimethoprim (stock solution at 50 mM diluted in 100 % dimethyl sulfoxide, DMSO). The mixture was incubated for 2 h 30 at room temperature (RT) before initiating crystallization experiments. Crystals were obtained in sitting drops at 18℃ using the Swissci MRC MAXI plates drops and by mixing 2 μL of protein solution with 2 μL of reservoir solution and equilibrated against a 200 μL reservoir solution made of 0.1 M Bis-Tris pH 6.5, 2.2 M AmSO4 and 2 % v/v Polyethylene glycol monomethyl ether 550. Prior to data collection, the crystals were mounted in a litholoop and cryocooled in liquid nitrogen.

### 6- Data collection and structure solving

The data collection was performed at the ID30B beamline at ESRF. 1400 frames were recorded with an oscillation range of 0.05°, an exposure time of 0.005 s, a beam transmission of 59%, a crystal-to-detector distance of 212.76 mm and at a wavelength of 0.873 Å on a EIGER 2 × 9M (Dectris). Data were processed with XDS [16] and the structure was solved by molecular replacement using the Phaser program [17] from the Phenix package [18] and using an AlphaFold v2 [19] model of the *N. asteroides* DHFR. The model was further adjusted in a series of cycle of rebuilding in Coot [20] and refinement with the Phenix package.

### 7- DHFR activity assays

The specific activities of WT DHFR and variants were assessed in a buffer composed of 50 mM Bis-Tris pH 5.5, 200 mM NaCl, 0.5 mg.mL^-1^ Bovine Serum Albumin (BSA) and 5 mM β-mercaptoethanol buffer. In brief, a 40 μL mix of the purified enzymes and NADPH at 40 nM and 2 mM respectively in solution with 2 % of DMSO for WT and M31A or 5 % for D30A and F34A, was first prepared and left for a 30 min incubation at RT in a 96-well plate (96 Well UV-Star® Microplates, Greiner). The reaction was started upon the addition of 40 µL of dihydrofolic acid (DHF, Sigma) at a concentration of 400 µM in the final mix. The absorbance at 340 nm, reflecting the consumption of NADPH thus the production of tetrahydrofolate (THF), was monitored over time in a Tecan Infinite 200 PRO M Plex plate reader at 25°C. The specific activity of three independent tests was calculated for each enzyme at 3 min and using a previously-established NADPH calibration curve containing 400 μM of DHF. Data and values were plotted on GraphPad Prism v10.6.1.

### 8- DHFR Inhibition assays

The half-maximal inhibitory concentration (IC_50_) was assessed for DMSO-solubilized trimethoprim and methotrexate (MTX) in the same buffer as previously described for the activity assays. To do so, a first mix of NADPH at 8 mM and DHFR at 160 nM was prepared by mixing 1:1 (v:v) the two solutions, and 20 µL of this mixture were transferred to two columns of a 96-well plate. 20 µL of a dilution of inhibitor ranging from 1 mM or 1 µM final of TMP or MTX, respectively, to 0 µM were added to these columns and left for incubation for 30 min at RT in the dark. The reaction was started upon the addition of 40 µL of a DHF solution at a concentration of 200 µM in the final reaction. The absorbance at 340 nm was monitored every 15 s for 45 s in a Tecan Infinite 200 PRO M Plex plate reader at 25°C. The A_t=0_ - A_t=45s_ was calculated and plotted on GraphPad Prism v10.6.1. The values were normalized according to the lowest and highest values obtained and the IC_50_ was retrieved for each test using the “log(inhibitor) vs. response – variable slope (four parameters)” analysis of the software. The final IC_50_ determined for each inhibitor is the result of the mean and standard deviation of three independent reactions. The same procedure was performed for the IC_50_ determination of TMP for the WT and two variants of DHFR (M31A and F34A), with the minor modifications that the DHF final concentration was increased to 400 µM, and the TMP dilutions ranged starting from 2.5 mM for F34A and from 1 mM for M31A. The final IC_50_ was calculated at 3 min for the WT and M31A enzymes, and at 20 min for the F34A variant.

### 9- Assessement of DHFR expression in *N. asteroides*

Cultures of 100 mL of *N. asteroides* transformed either with pNV118_Zeo_::*dhfr*_*Nad*_ or pNV118_Zeo_::*eGFP* were grown exponentially (2 to 5 days) in liquid BHI supplemented with 50 μg.mL^-1^ of zeocin at 30℃. The cultures were centrifuged for 10 min at 3500 *g* at RT. The pellet was washed with 25 mL of Phosphate-Buffered Saline (PBS) 1X and spun down for 10 min. The pellet was resuspended in 6 mL of PBS 1X supplemented with 10 % glycerol and 2 mL of beads (0.1 mm silica spheres, lysing matrix B, PM Biomedicals) and transferred into a 50 mL falcon tube. The samples were then shaken for 45 s at a speed of 6 m.s^-1^. This procedure was repeated 4 times with cooling in ice for several minutes between each cycle. The extract was then centrifuged for 10 min at 3500 *g* and the supernatant was spun down for 10 min at 15000 *g*.

For protein enrichment by affinity chromatography, the extracts were then loaded onto a gravity column containing about 500 μL of Strep-Tactin™ Superflow™ High-Capacity Resin (IBA Life-sciences) equilibrated with PBS 1X and 10 % glycerol buffer. The flow-through was reloaded 5 times to ensure proper binding. The columns were then washed with 10 volumes of buffer. Proteins were eluted by incubating the beads for 30 min with 500 µL of buffer containing 2.5 mM of desthiobiotin. To assess DHFR expression, proteins from either the crude extract or from the eluted fraction from the Strep-Tactin beads were separated by a 15 % SDS-PAGE gel. Proteins were then transferred onto Nitrocellulose Blotting membranes (Amersham™ Protran™ 0.2 μm NC) using a Bio-Rad liquid transfer apparatus at a constant voltage of 100 V for 1 h and using the following transfer buffer: 24 mM Tris, 192 mM glycine and 20 % ethanol. Membranes were then blocked in 1X PBS and 10 % (w/v) milk for 30 min at RT and further incubated ON at 4 °C with a monoclonal anti-Strep-Tag II primary antibody (Sigma, 1:2000 dilution) in PBS-Tween 0.1 % containing 5 % (w/v) milk. Membranes were washed four times for 10 min each in PBS-Tween 0.1 %, followed by incubation with an HRP-conjugated anti-mouse secondary antibody (Sigma, 1:2000 dilution) for 3 h at RT. After four additional washes in PBS-Tween 0.1 %, the signal was detected using the Clarity Max™ Western ECL substrate (Bio-Rad).

### 10- Antimicrobial susceptibility testing

*N. asteroides* strains were grown into liquid BHI media supplemented with zeocin (50 μg.mL^-1^) at 30 ℃ under agitation. The bacteria were then pelleted by centrifugation and resuspended in sterile Phosphate Buffer Saline 1X (PBS 1X) followed by a new centrifugation step. The pellet was then resuspended in PBS 1X. Bacterial aggregates were dissociated by 20 passaging into a 1 mL tuberculin syringe plugged with a G26 needle. Once the bacterial solution was homogeneous the OD_600nm_ was adjusted to 1 and a 1:10 serial dilution series was performed in PBS 1X. 2 μl of the bacterial dilutions were spotted on Mueller-Hinton agar plates supplemented or not with TMP at concentrations ranging from 50 to 1.6 μg.mL^-1^. Plates were placed at 30℃ observed and photographed at 96 h.

### 11- Multiple sequence alignment and structural analysis

DHFR protein sequences were retrieved from the KEGG (https://www.genome.jp/kegg/) or UniProt (https://www.uniprot.org/) servers. Multiple sequence alignment was performed on the Clustal Omega server (https://www.ebi.ac.uk/jdispatcher/msa/clustalo). Structural alignment was performed on the ESPript server [21] and structure superpositions and analysis were performed with PyMOL version 2.5.0, Schrödinger, LLC.

## Results and Discussion

### Overexpression of DHFR_Nad_ in *N. asteroides* triggers trimethoprim resistance

DHFR overexpression as a mechanism of TMP resistance has been known for several decades. It was reported in a highly resistant *E. coli* clinical strain that a mutation in the *folA* promoter region inducing a massive expression of the DHFR triggers TMP resistance [22]. This was also confirmed in an *E. coli* strain that overexpressing DHFR artificially enhances resistance to TMP [23].

To identify and confirm the drug target of TMP in *N. asteroides*, we opted for an overexpression approach. First, to address and identify the DHFR encoding gene from *N. asteroides*, we ran a BLAST search on the UniProt database against its genome and using the *Mycobacterium tuberculosis* H37Rv strain DHFR protein sequence (Rv2763c) as a search request.

We identified the UniProt entry U5E929 as the best match which corresponds to an open reading frame of 162 amino acids sharing 56 % protein sequence identity with the DHFR from *M. tuberculosis*. As no other significant hit was found, we amplified by PCR the corresponding gene from the genomic DNA of *N. asteroides* and cloned it into the pNV118 plasmid. We designed the construct so that a Strep-Tag II peptide is present at the C-terminus of the expressed protein enabling its purification onto an affinity chromatography column and assessment of its expression by Western-Blot (WB). The resulting construct pNV118_Zeo_::*dhfr*_*Nad*_ was electroporated into *N. asteroides* strain. As a negative control, we cloned the gene encoding for eGFP between the same cloning sites and plasmid backbone. Using this design, we could detect the protein in a crude extract of *N. asteroides* by WB and using an anti-Strep-Tag II primary antibody (**FIGURE 1A)**. A band corresponding to the molecular weight of DHFR (∼18 kDa) could be detected in the clone transformed with pNV118_Zeo_::*dhfr*_*Nad*_ but not in the one transformed with pNV118_Zeo_::*eGFP*. Further, in another independent experiment, we also try to enrich the protein by loading the supernatant after bacterial lysis over a Strep-Tactin column. After elution, the presence of DHFR was also assessed similarly by WB. This approach also confirmed the first results obtained from the crude extract (**FIGURE S1A**, supplementary materials). Altogether, these data attested of the expression of DHFR in the clone transformed with pNV118_Zeo_::*dhfr*_*Nad*_ but rather weak expression of DHFR as we did not notice a clear enrichment of the protein after affinity chromatography.

**FIGURE 1.**
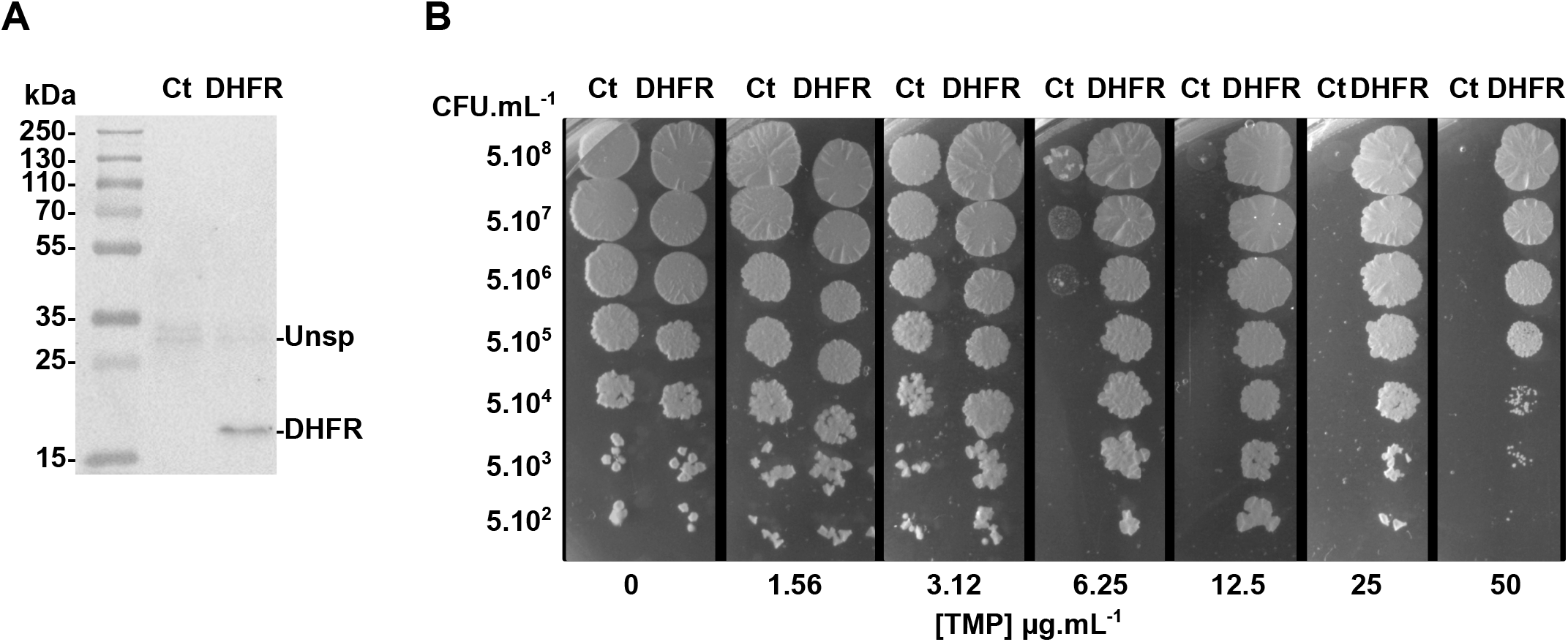
DHFR overexpression in *N. asteroides* induces drug resistance to trimethoprim. **A-** The expression of DHFR fused to Strep-Tag II was estimated in a crude extract of *N. asteroides* transformed with pNV118_Zeo_::*dhfr*_*Nad*_. A specific band at about 18 kDa was detected (DHFR) but was not present in the control strain transformed with pNV118_Zeo_::*eGFP* (Ct). An unspecific and non-identified band at about 30 kDa was seen in both extracts (Unsp). **B-** Drug susceptibility testing of *N. asteroides* to trimethoprim. The susceptibility of the two strains transformed with either pNV118_Zeo_::*eGFP* (Ct) or pNV118_Zeo_::*dhfr*_*Nad*_ (DHFR) were assessed in the absence or presence of increased concentrations of trimethoprim (TMP) ranging from 1.5 to 50 μg.mL^-1^. Three biological and independent experiments were performed. The picture is a representative of one of them, the two other experiments are shown in supplementary **FIGURE S1B and S1C**.

We next assessed and compared the susceptibility to TMP of the two strains on Mueller-Hinton agar plate with increasing concentrations of TMP from 0 to 50 μg.mL^-1^ and a serial dilution of *N. asteroides* **(FIGURES 1B and S1B**,**C)**. First of all, *N. asteroides* strains overexpressing DHFR and eGFP (Ct) grew similarly on Mueller-Hinton agar plates without antibiotic.The susceptibility of the two strains to TMP was barely affected at 1.5 μg.mL^-1^ of TMP but a growth defect appeared at 3.1 μg.mL^-1^ for the control strain expressing eGFP while the DHFR expressing strain was not affected. The control strains did not grow anymore at 6.2 μg.mL^-1^ while the strain expressing DHFR could still grow at 50 μg.mL^-1^ except for the highest dilution **(FIGURE 1B and S1B**,**C)**.

These data clearly demonstrated that overexpression of DHFR triggered a high degree of TMP resistance and strongly confirmed the identified gene as the one encoding for DHFR. Thus, it was renamed DHFR_Nad_.

### Biochemical characterization of DHFR_Nad_ and its inhibition by TMP

To further confirm the data from the overexpression approach, we engaged in biochemical characterization of DHFR_Nad_. For this purpose, a synthetic construct of codon-optimized DHFR_Nad_ was recombinantly expressed in *E. coli* and purified via a three-step purification procedure. After affinity purification on IMAC, cleavage and removal of the SUMO tag, the protein behaved as a very homogeneous monomer peak as demonstrated by the SEC elution profile. DHFR_Nad_ was very pure after this purification protocol **(FIGURE 2)**. We then assessed the protein activity by demonstrating its capacity to produce tetrahydrofolate from dihydrofolate in the presence of NADPH. As shown in **FIGURE 3A**, we could indeed detect such activity at slightly acidic pH.

**FIGURE 2.**
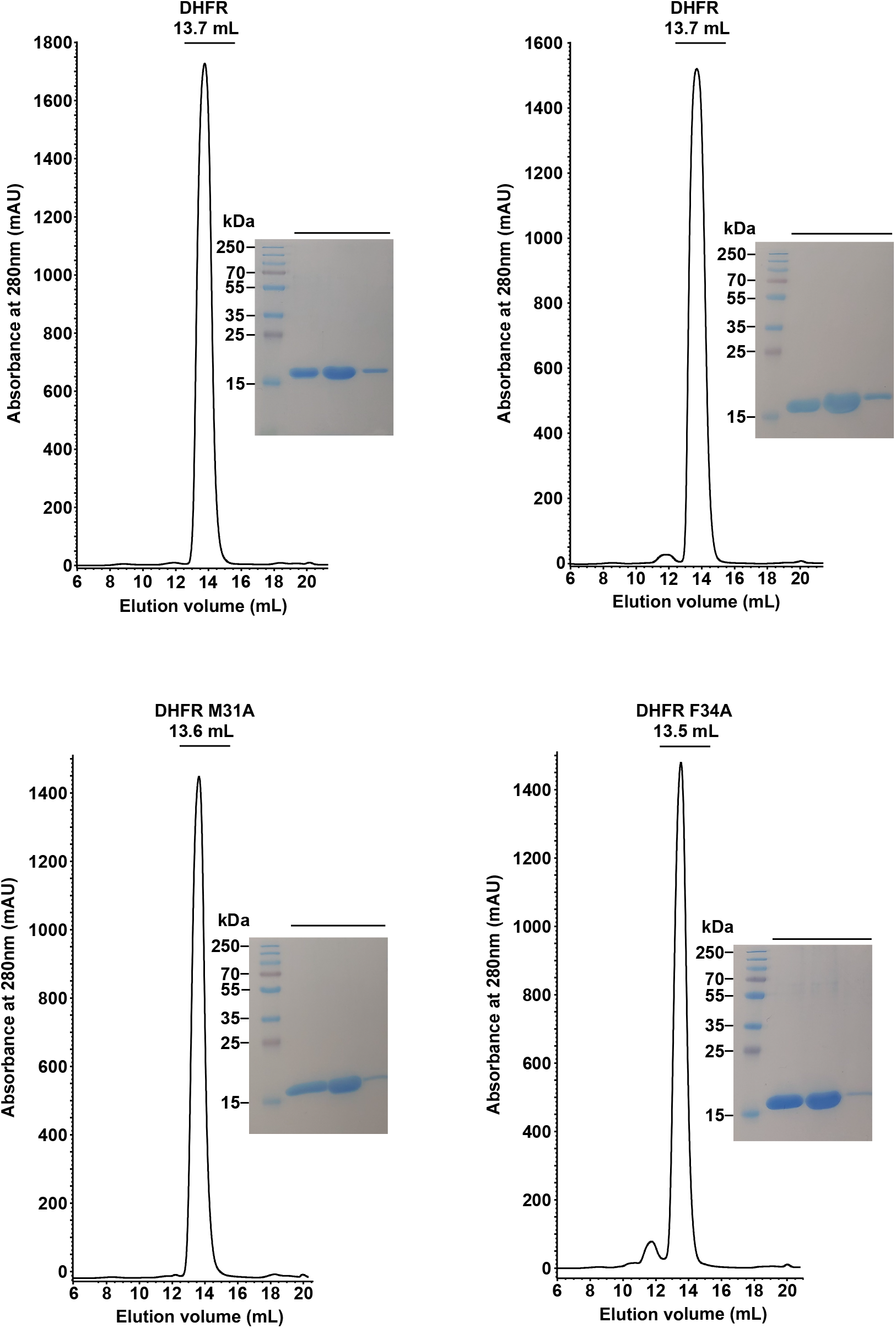
Purification of WT DHFR and variants recombinantly expressed in *E. coli*. The figure depicts the chromatograms from the last step of purification by size-exclusion chromatography on a Superdex 75 Increase 10/300 GL column. Three fractions from the elution peak were pooled for each mutant and their purity was analyzed on Coomassie blue-stained SDS-PAGE. All variants eluted at similar elution volumes and behaved as monodispersed and homogeneous peaks, suggesting that the single point mutations did not trigger any folding issue.

**FIGURE 3.**
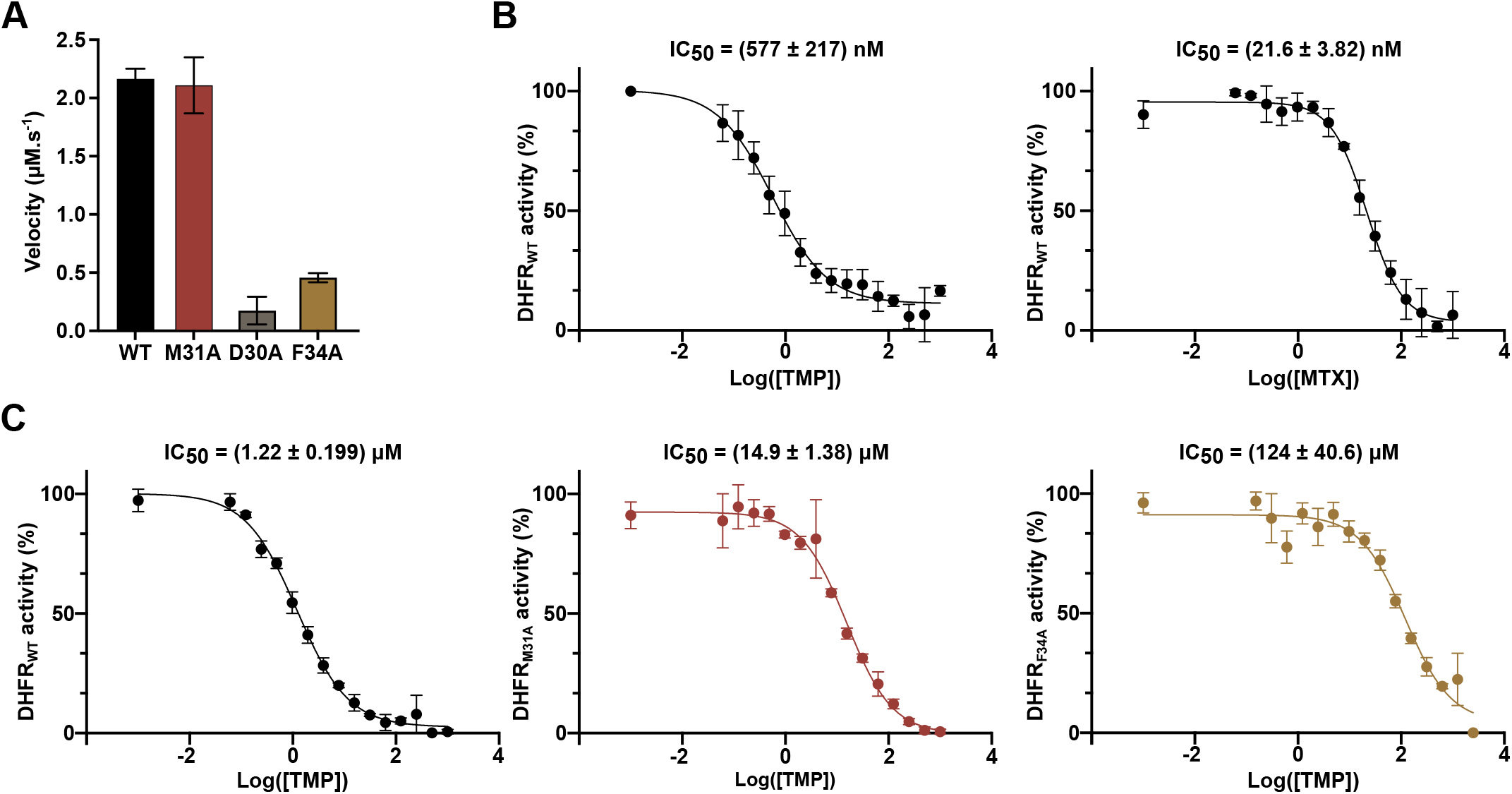
*In vitro* DHFR activity assays. **A-** Determination of the WT and variants activities of DHFR_Nad_. The initial velocity of each variant was assessed and calculated after 3 min of reaction. **B-** Inhibition of DHFR_Nad_ by TMP and MTX. The IC_50_ were determined under similar conditions and with 200 μM of substrate. **C-** Inhibition of DHFR_Nad_ WT and variants by TMP. The IC_50_ were determined under similar conditions and with 400 μM of substrate. All IC_50_ values correspond to the mean of three independent experiments and the ± indicates the standard deviation.

We next confirmed that the protein of interest is the target of TMP by testing its activity in the presence of increasing concentrations of TMP, and further determined the IC_50_ of the drug. We found an IC_50_ of 577 nM which is much higher than for *E. coli* DHFR that has an IC_50_ of 20 nM [24] however much lower than the one of the closely related *M. tuberculosis* or *Mycobacterium abscessus* with IC_50_ of 19 μM [25] or 25 μM [26] respectively **(FIGURE 3B)**.

We determined as well the IC_50_ for another inhibitor methotrexate (MTX) that exhibited more than 20 times higher inhibition properties than TMP with an IC_50_ of 21 nM **(FIGURE 3B)**. This is consistent with Mycobacterial DHFR which are more inhibited by MTX than with TMP with an IC_50_ of 6.7 nM for *M. tuberculosis* DHFR and an IC_50_ of 13 nM for that of *M. abscessus*. Albeit not used as an antimicrobial drug, the fact that MTX is harboring a much better IC_50_ for DHFR_Nad_ demonstrates the possibility to find more potent Nocardia DHFR inhibitors.

### Crystal structure of DHFR_Nad_ bound to NADPH and trimethoprim

We next supported these biochemical data by obtaining the crystal structure of DHFR_Nad_ **(FIGURE 4A)**. The DHFR_Nad_ structure was solved and refined at a resolution of 1.55 Å **(Table I)**. We could rebuild all the residues 2-162 that were present in the construct after cleavage of the affinity and solubility tags. Additionally, 2 sulfate ions were placed and a clear electron density map was visible for two extra ligands present in the co-crystallization experiment, namely NADPH and trimethoprim, as judged by the Fo-Fc SA-OMIT map **(FIGURE 4B)**.

**Table I:**
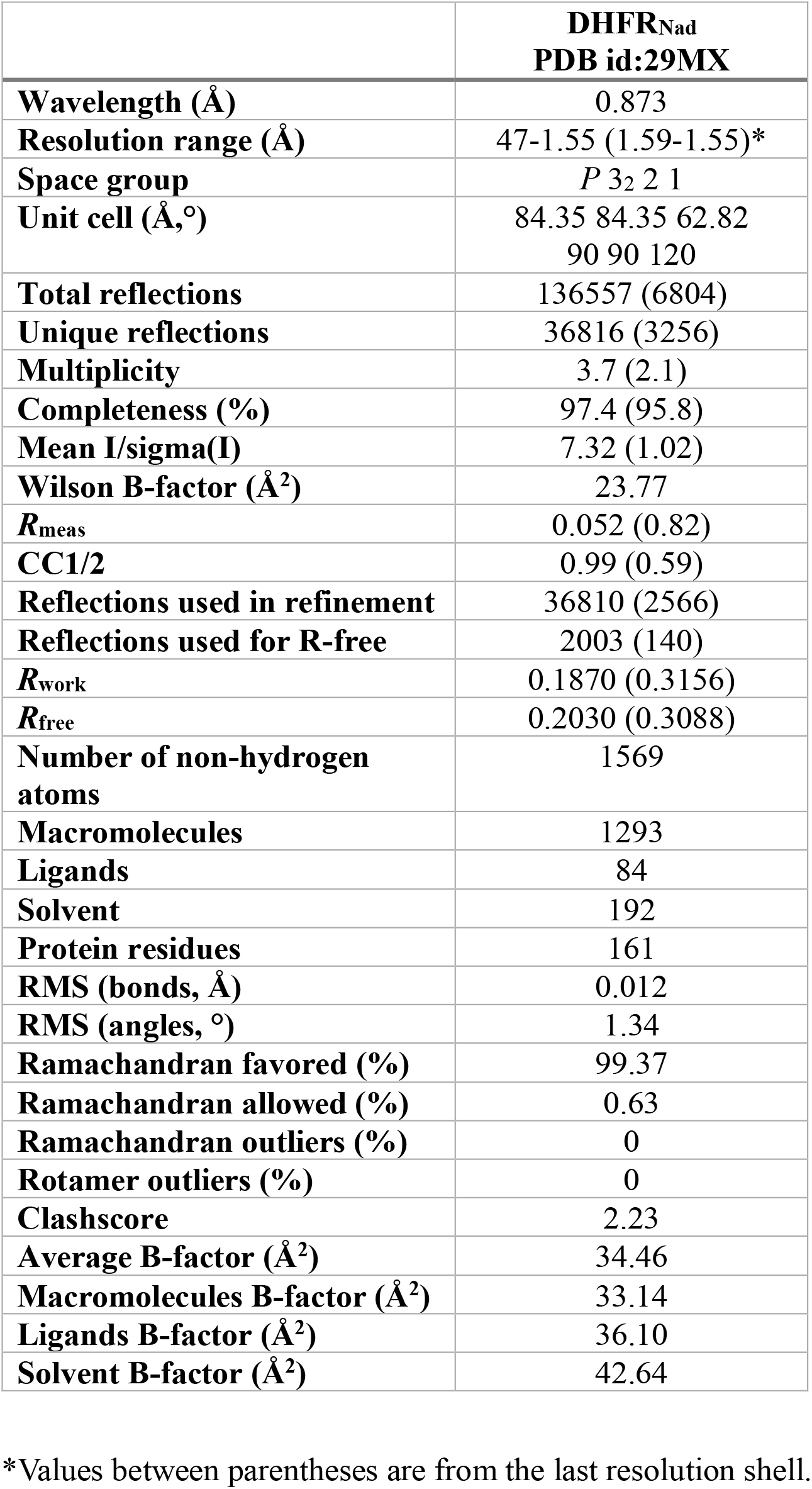
X-ray data collection and refinement statistics

|  | **DHFR<sub>Nad</sub>**<br>**PDB id:29MX** |
| --- | --- |
| **Wavelength (Å)** | 0.873 |
| **Resolution range (Å)** | 47-1.55 (1.59-1.55)* |
| **Space group** | *P* 3<sub>2</sub> 2 1 |
| **Unit cell (Å,°)** | 84.35 84.35 62.82<br>90 90 120 |
| **Total reflections** | 136557 (6804) |
| **Unique reflections** | 36816 (3256) |
| **Multiplicity** | 3.7 (2.1) |
| **Completeness (%)** | 97.4 (95.8) |
| **Mean I/sigma(I)** | 7.32 (1.02) |
| **Wilson B-factor (Å<sup>2</sup>)** | 23.77 |
| **R<sub>meas</sub>** | 0.052 (0.82) |
| **CC1/2** | 0.99 (0.59) |
| **Reflections used in refinement** | 36810 (2566) |
| **Reflections used for R-free** | 2003 (140) |
| **R<sub>work</sub>** | 0.1870 (0.3156) |
| **R<sub>free</sub>** | 0.2030 (0.3088) |
| **Number of non-hydrogen atoms** | 1569 |
| **Macromolecules** | 1293 |
| **Ligands** | 84 |
| **Solvent** | 192 |
| **Protein residues** | 161 |
| **RMS (bonds, Å)** | 0.012 |
| **RMS (angles, °)** | 1.34 |
| **Ramachandran favored (%)** | 99.37 |
| **Ramachandran allowed (%)** | 0.63 |
| **Ramachandran outliers (%)** | 0 |
| **Rotamer outliers (%)** | 0 |
| **Clashscore** | 2.23 |
| **Average B-factor (Å<sup>2</sup>)** | 34.46 |
| **Macromolecules B-factor (Å<sup>2</sup>)** | 33.14 |
| **Ligands B-factor (Å<sup>2</sup>)** | 36.10 |
| **Solvent B-factor (Å<sup>2</sup>)** | 42.64 |

\*Values between parentheses are from the last resolution shell.

**FIGURE 4.**
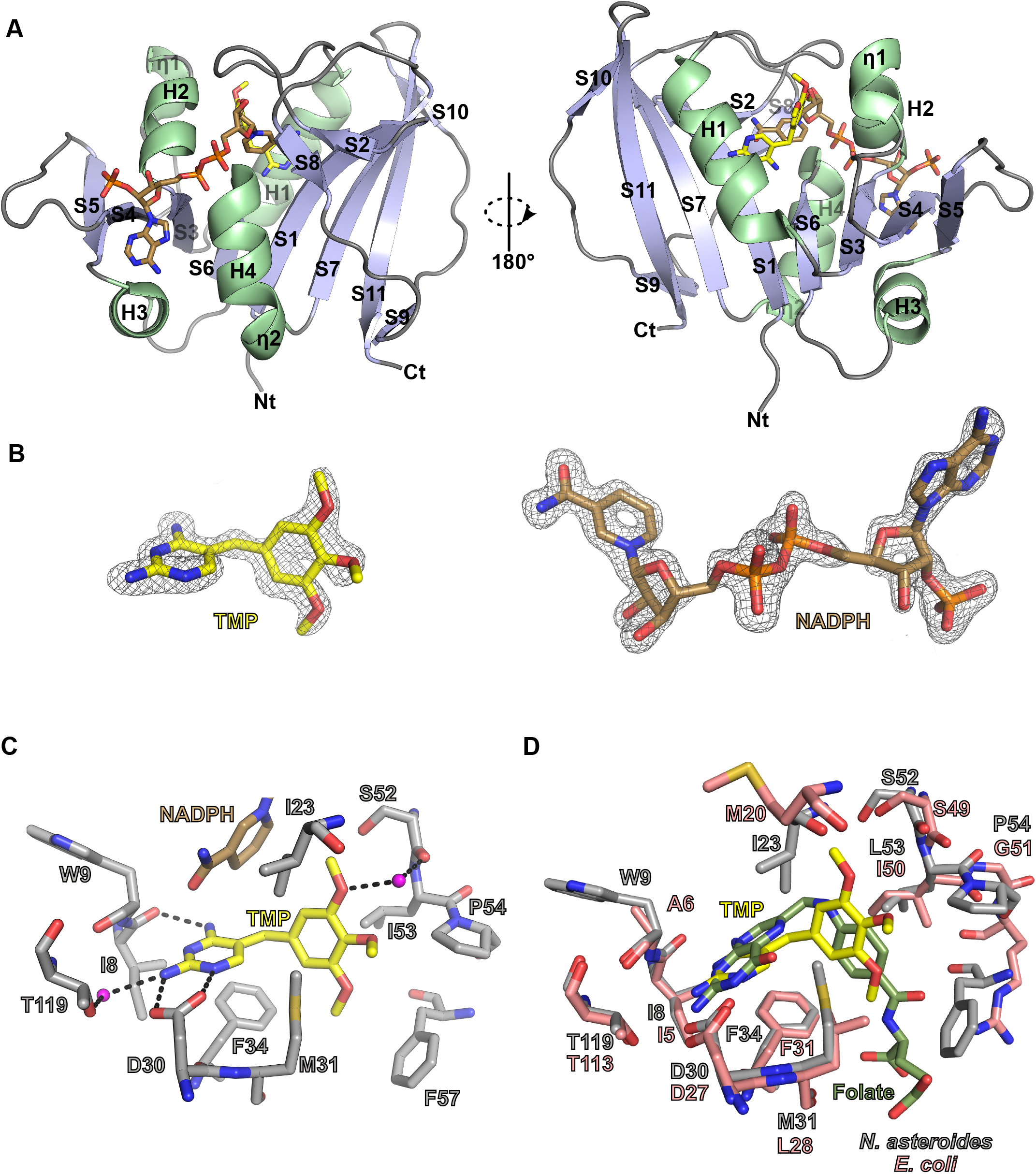
Structural analysis of DHFR_Nad_. **A-** Overall structure of DHFR_Nad_. The structure is shown as a cartoon representation in two different orientations. β-strands (S1 to S11) are in light blue, while ⍺-helices (H1 to H4) and 3_10_ helices are in light green. Coils are in gray. The two ligands, trimethoprim and NADPH, are represented as yellow and brown sticks, respectively. **B-** Simulated annealed OMIT map contoured at 3.5 σ around each ligand and shown as a grid mesh. **C-** Zoom on the binding site of TMP. Dash lines indicate H-bond interactions and the pink spheres represent water molecules. TMP is represented in yellow sticks, while for the NADPH cofactor only the nicotinamide group is represented as sticks and colored in brown. **D-** Superposition of the *E. coli* (in pink) (PDB id: 6R2A) and *N. asteroides* (in grey) DHFR structures.

The overall DHFR_Nad_ structure is made of a central β-sheet surrounded by four ⍺-helices and is very similar to DHFR structures from other phylae **(FIGURE 4A)**. A search in the PDB using the PDBeFold server (https://www.ebi.ac.uk/msd-srv/ssm/) retrieves as the closest DHFR structures the ones from *Mycobacterium* species. The nearest structure is the one from *Mycobacterium abscessus* bound to NADP and an inhibitor (PDB 7k6c) [27]. The two structures share indeed a r.m.s.d. of 0.86 Å over 157 residues on their carbon alpha. This high similarity was expected as *Nocardia* and *Mycobacteria* are closely related bacteria belonging to the same order.

The TMP inhibitor is situated in proximity to the nicotinamide group of NADPH and is tightly bound by several residues **(FIGURE 4C)**. The main chain of I8, via its carbonyl group establishes a H-bond with the N7 atom of the pyrimidine moiety, which is also recognized via a salt-bridge interaction with the side chain of D30. The pyrimidine group is recognized by F34 mediating a stacking interaction and a H-bond via a water molecule by the side chain of T119. The 1,2,3-methoxybenzene moiety of TMP is contacted by a hydrophobic interaction with the side chain of I23 and by a hydrogen bond via a water molecule with the main chain of S52. Finally, the binding pocket is closed with the side chains of M31, L53, P54 and F57. Comparison of the structure with the one of *E. coli* DHFR bound to NADP and folate (PDB 1RA2) [28] clearly attests as well and as expected that TMP is binding in the same cavity as the DHFR substrate **(FIGURE 4D)**.

As this study is the first of its kind for the *Nocardia* field of research and in the aim to extrapolate the data to other commonly studied *Nocardia* species, we performed sequences alignment of DHFR_Nad_ with those of five other *Nocardia* species. This comparison attested that DHFR_Nad_ shares 64 % sequence identity with the DHFR from *N. farcinica*, 62 % with that of of *N. nova*, 70 % with that of *N. otitidiscaviarum*, 71 % with those of *N. brasiliensis* and *N. cyriacigeorgica*. This analysis thus demonstrated that the DHFR_Nad_ structure is a good representative of other DHFR *Nocardia* species **(FIGURE 5)**.

**FIGURE 5.**
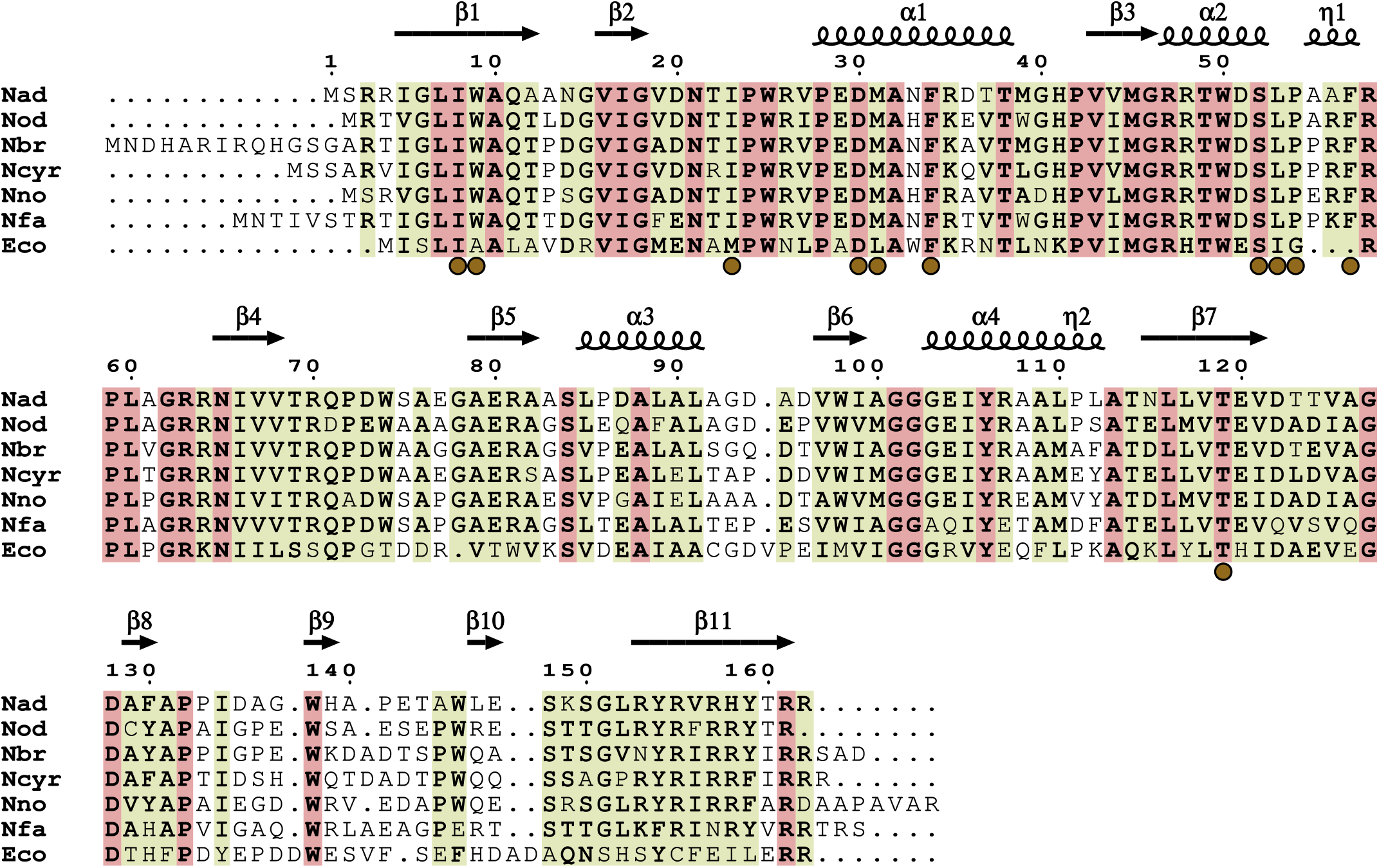
Multiple sequence alignment. The sequence of DHFR_Nad_ (Nad) was aligned with the sequences of DHFR from *Nocardia otitidiscaviarum* NEB252 (Nod), *Nocardia farcinica* IFM 10152 (Nfa), *Nocardia brasiliensis* ATCC 700358 (Nbr), *Nocardia nova* SH22a (Nno) and *Nocardia cyriacigeorgica* GUH-2 (Ncyr). The *E. coli* (Eco) DHFR sequence was added as it is a well-studied and a reference enzyme in the DHFR field of research. Strictly-conserved residues are in light red while semi-conserved ones are in light green. The brown spheres below the sequences indicate the residues involved in TMP recognition. The secondary structure elements of the DHFR_Nad_ structure are placed above the sequence alignment where α, β, η indicates α-helix, β-sheet and 3_10_ helix respectively. The figure was generated with the ENDscript server [32] and manually adjusted in Inkscape.

### Mutagenesis analysis highlights key residues for DHFR_Nad_ activity, TMP binding and drug resistance

To validate the binding pose of TMP seen in the co-crystal structure, we generated and purified three variants of DHFR_Nad_ namely the D30A, the M31A and F34A versions, that are all involved in the binding of TMP. We could express and purify these variants following the same procedure as for the wild-type (WT) enzyme. They all eluted at the same volume as the WT version on the SEC column suggesting that the mutations did not affect their folding **(FIGURE 2)**. We next tested the activity of all the mutants in comparison with the WT and could show that while the M31A variant is as active as the WT, the two others had a reduced activity **(FIGURE 3A)**. This observation was somehow expected for the D30 residue substitution as a previous study on *E. coli* DHFR demonstrated the equivalent residue (D27) **(FIGURE 4D)** to be critical for the activity of the enzyme [29] and in line with the fact that it is involved in the recognition of the folate substrate [30]. It was also shown that the substitution of the F34 equivalent residue in the *Pneumocystis jirovecii* DHFR reduced its activity as well [31]. This F residue is also involved in substrate recognition as seen in the *E. coli* co-crystal structure **(FIGURE 4D)**.

We next determined the IC_50_ of TMP for these enzyme variants. We could not determine the IC_50_ for the D30A variant as its activity was too weak **(FIGURE 3A)**. However, we could show that the M31A and F34A are less susceptible to TMP than the WT with IC_50_ values of 14.9 μM and 124 μM, respectively, corresponding to 12- and 101-fold increases as compared to the WT (1.2 μM) in similar conditions **(FIGURE 3C)**. It is important to note that for these experiments, we had to increase the substrate concentration to 400 μM to compensate for the weaker activity of the F34A variant. This also explains why the IC_50_ for WT is different in **FIGURE 3B** and **C**. Altogether, these results validated the role of these residues in the binding of TMP. Interestingly, the low sensitivity of the M31 variant to TMP is in line with a previous study that demonstrated that some spontaneously TMP drug-resistant *Nocardia cyriacigeorgica* mutants displayed a mutation in the equivalent M31 residue [10]. Indeed, these M30R *N. cyriacigeorgica* mutants displayed very high MIC ranging from 62 to 608 μg.mL^-1^ to cotrimoxazole. Despite resistance to TMP was not solely evaluated in that study, it nonetheless highlighted this methionine residue to be crucial for resistance mechanism to anti-nocardiosis treatment, as shown in our biochemical studies.

## Conclusion

DHFR and DHPS are the two main targets of the anti-nocardiosis treatment. Before this work, no study had demonstrated the direct inhibition of these enzymes by trimethoprim or sulfamethoxazole. This study characterized the DHFR from *N. asteroides* which is one of the most frequently encountered species in Nocardiosis. We were able to purify and characterize the enzyme and demonstrate that TMP is a potent inhibitor for IC_50_ in the ηM or low μM range. Our structural work demonstrates that the *Nocardia* DHFR is closely related to other bacterial DHFRs and particularly those of mycobacteria. Furthermore because of the high sequence similarity with other DHFR from pathogenic *Nocardia* species our work could be of interest to interpret mutations in resistant strains to first line treatment found in the clinic. Finally, the fact that methotrexate harbors a much better IC_50_ than that of TMP offers the possibility to find more potent inhibitors of Nocardia DHFR and that could be of great interest, as side effects of first line treatment are frequent.

## Acknowledgements

We thank the Leibniz Institute DSMZ-German Collection of Microorganisms and Cell Cultures GmbH, for providing the *N. asteroides* strain and Dr. J. Ishikawa for kindly sharing the pNV118 plasmid. We acknowledge the European Synchrotron Radiation Facility (ESRF) for provision of synchrotron radiation facilities. We would like to thank the staff of the ESRF Grenoble for assistance and support in using beamline ID30B under proposal number MX-2600. This project was funded by ANR-24-CE44-5883.

## CRediT

**Julie Couston:** Conceptualization, Methodology, Formal analysis, Investigation, Writing – review and editing. **Sébastien Lainé:** Conceptualization, Methodology, Formal analysis, Investigation, Writing – review and editing. **Jérôme Feuillard:** Formal analysis, Investigation, Writing – review and editing. **Mickaël Blaise:** Conceptualization, Methodology, Formal analysis, Investigation, Resources, Project administration, Validation, Supervision, Writing – original draft, Writing – review and editing, Funding acquisition.

## Data availability

Data relative to X-ray diffraction and collected under the ESRF proposal ID: MX-2600 are available in the following repositories: https://doi.org/10.15151/ESRF-DC-2432612030. The final structure coordinates and structure factors were deposited at the Protein Data Bank under the following accession number: 29MX. The other data that support the findings of this study are available from the corresponding authors upon reasonable request.

