## Supplementary FigureS1 for "Biochemical, structural, and functional characterization of the *Nocardia asteroides* dihydrofolate reductase: a primary target of anti-nocardiosis treatment"

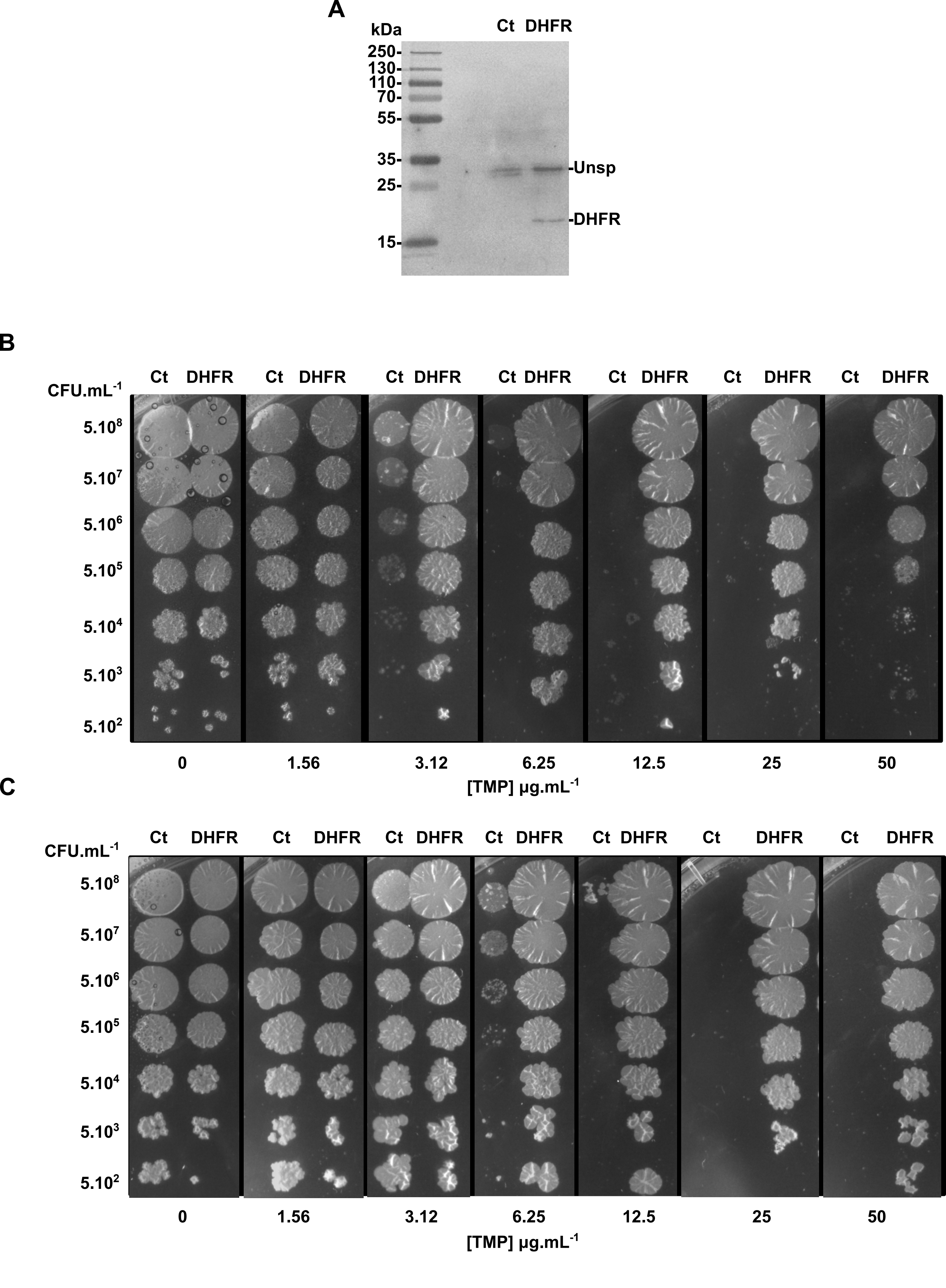
A
kDa
Ct DHFR
250-
130-
110-
70-
55-
35-
25-
15-
-Unsp
-DHFR
B
Ct DHFR Ct DHFR Ct DHFR Ct DHFR Ct DHFR Ct DHFR Ct DHFR
CFU.mL-1
5.10^8
5.10^7
5.10^6
5.10^5
5.10^4
5.10^3
5.10^2
0 1.56 3.12 6.25 12.5 25 50
[TMP] µg.mL-1
C
Ct DHFR Ct DHFR Ct DHFR Ct DHFR Ct DHFR Ct DHFR Ct DHFR
CFU.mL-1
5.10^8
5.10^7
5.10^6
5.10^5
5.10^4
5.10^3
5.10^2
0 1.56 3.12 6.25 12.5 25 50
[TMP] µg.mL-1

**Figure S1**

**Figure Supplementary S1: DHFR overexpression in *N. asteroides* induces drug resistance to trimethoprim**

**A-**The expression of the DHFR fused to Strep-tag II was estimated after an affinity chromatography step. A specific band at about 18 kDa was detected (DHFR) but was not present in the control strain transformed with pNV118$_{Zeo}$::*eGFP* (Ct). An unspecific and non-identified band at about 30 kDa was seen in both extracts. **B-** and **C-** Drug susceptibility testing of *N. asteroides* to trimethoprim. The susceptibility of the two strains transformed with either pNV118$_{Zeo}$::*dhfr*$_{Nad}$ (DHFR) or pNV118$_{Zeo}$::*eGFP* (Ct) were assessed in the absence or presence of increased concentrations of trimethoprim ranging from 1.5 to 50 µg.mL$^{-1}$. The picture displayed the two replicates performed in addition to the experiment presented in the main Figure 1.
